# Tetraploidy-linked sensitization to CENP-E inhibition in human cells

**DOI:** 10.1101/2022.08.21.504625

**Authors:** Koya Yoshizawa, Akira Matsura, Masaya Shimada, Sumire Ishida-Ishihara, Takahiro Yamamoto, Kan Yaguchi, Eiji Kawamoto, Taruho Kuroda, Kazuya Matsuo, Nobuyuki Tamaoki, Ryuichi Sakai, Mithilesh Mishra, Ryota Uehara

**Affiliations:** Graduate School of Life Science, Hokkaido University, Japan; Faculty of Advanced Life Science, Hokkaido University, Japan; Graduate School of Medicine, Mie University, Japan; Faculty of Molecular Chemistry and Engineering, Kyoto Institute of Technology, Kyoto, Japan; Research Institute for Electronic Science, Hokkaido University, Japan; Graduate School and Faculty of Fisheries Sciences, Hokkaido University, Japan; Department of Biological Sciences, Tata Institute of Fundamental Research, India

## Abstract

Tetraploidy caused by whole-genome duplication is a hallmark of cancer cells, and tetraploidy-selective cell growth suppression is a potential strategy for targeted cancer therapy. However, how tetraploid cells differ from normal diploids in their sensitivity to anti-proliferative treatments remains largely unknown. In this study, we found that tetraploid cells are significantly more susceptible to inhibitors of a mitotic kinesin CENP-E than diploids. CENP-E inhibitor preferentially diminished the tetraploid cell population in diploid-tetraploid co-culture at optimum conditions. Live imaging revealed that tetraploidy-linked increase in unsolvable polar chromosome misalignment caused substantially longer mitotic delay in tetraploids than in diploids upon moderate CENP-E inhibition. This time gap of mitotic arrest resulted in cohesion fatigue and subsequent cell death, specifically in tetraploids, leading to tetraploidy-selective cell growth suppression. In contrast, the microtubule-stabilizing compound paclitaxel caused tetraploidy-selective growth suppression through the aggravation of spindle multipolarization. We also found that CENP-E inhibitor had superior generality to paclitaxel in its tetraploidy selectivity across a broader spectrum of cell lines. Our results highlight the unique properties of CENP-E inhibitors in tetraploidy-selective suppression, giving us clues on the further development of tetraploidy-targeting interventions in cancer.

## Introduction

Tetraploidy resulting from whole-genome duplication (WGD) of a normal diploid cell is a common hallmark of cancer. Recent cancer genome analyses revealed that about 30% of solid tumors had undergone at least one round of WGD ^1, 2^. The induction of tetraploidization facilitates tumorigenesis and malignant transformation in mice models, suggesting that tetraploidy is a critical intermediate state in these pathogenic processes ^3, 4^. The principle of tetraploidy-driven cancer formation is still largely unknown. However, recent studies have proposed that increased tolerance to chromosome alterations and instability or enhanced invasiveness upon tetraploidization contribute to the oncogenic quality of tetraploid cells ^5–7^. Because of the commonality and significant contributions of tetraploidy to the tumorigenic process, selective suppression of tetraploid cell growth is a promising strategy for cancer chemotherapy ^8^. In this context, mitosis is a good candidate for the tetraploidy-selective chemotherapeutic target. A previous study reported that tetraploid hTERT-RPE1 cells took longer to go through mitosis than diploid counterparts even when they had the normal number (i.e., 2) of centrosomes ^9^, suggesting that the doubled number of chromosomes increases the burden on the mitotic mechanism upon tetraploidization.

Moreover, recent studies revealed that tetraploid cells are more susceptible to anti-mitotic microtubule stabilizer paclitaxel or inhibitors of a mitotic kinase MPS1, Plk1, or a mitotic kinesin motor protein Kif18A ^10–13^. These findings suggest that tetraploid cells have an increased dependence on specific aspects of mitotic regulations, presumably as adaptive mechanisms to the increased burden of doubled chromosomes. Elucidation of such tetraploidy-linked adaptive mechanisms would provide more choices of tetraploidy-selective cell growth suppression, potentially benefiting the development of tetraploidy-targeting chemotherapeutic strategy in broad cancer types.

Centromere-associated protein E (CENP-E; kinesin-7) is a mitotic kinesin that plays an essential role in transporting mitotic chromosomes along spindle microtubules and aligning them on the equatorial metaphase plate ^14–16^. Inhibition of CENP-E’s ATPase activity by an allosteric inhibitor GSK-923295 causes tight binding of the protein to microtubules, resulting in frequent chromosome misalignment at the spindle poles and mitotic arrest through the activation of the spindle assembly checkpoint (SAC) ^14, 17, 18^. The specific requirement of CENP-E in mitosis makes it an ideal candidate for an anti-mitotic cancer therapeutic target ^19, 20^. In mitosis, not all chromosomes require CENP-E activity for their alignment. While the large population of mitotic chromosomes can align at the equatorial plate, those initially located in the nuclear peripheral region upon mitotic entry tend to be trapped at the spindle pole in the absence of CENP-E activity ^21^. Moreover, while smaller-sized chromosomes tend to re-align to the equatorial plate even when initially trapped at the spindle poles, larger-sized chromosomes have less chance of re-alignment ^22^. Therefore, the location and size of the mitotic chromosomes affect their susceptibility to CENP-E inhibition. On the other hand, it remains unclear whether and how drastic differences in chromosome number affect cellular susceptibility to CENP-E inhibition.

In this study, we compared the effect of anti-mitotic compounds on the proliferation of cells at different ploidy states. Among these compounds, CENP-E inhibitors significantly suppressed the proliferation of tetraploid cells compared to diploids in different culture conditions or cellular backgrounds. We found that the tetraploidy-selective suppression was based on the aggravation of chromosome misalignment, mitotic arrest, and consequent cell death upon CENP-E inhibition. On the other hand, paclitaxel caused tetraploidy-selective cell death via the aggravation of mitotic spindle multipolarization, highlighting the difference in the principle of tetraploidy-selective cell growth suppression by paclitaxel and CENP-E inhibitors. We also found that a CENP-E inhibitor showed selectivity toward a broader spectrum of tetraploid cell lines compared to paclitaxel, demonstrating superior generality of CENP-E-targeted tetraploidy suppression. Based on our results, we discuss the potential values of various tetraploidy-targeting mechanisms of different anti-mitotic compounds.

## Results

### Selective suppression of tetraploid cell growth by CENP-E inhibitors

To understand the influence of ploidy difference on cellular sensitivity to mitotic perturbations, we compared the effect of various anti-mitotic compounds on isogenic haploid, diploid, and tetraploid HAP1 cells ^23^ (Fig. S1A) using a colorimetric cell proliferation assay. Different compounds showed diverse trends and varying degrees of ploidy dependency in efficacy (Fig. 1A, S2, and S3). Therefore, we categorized these compounds based on statistical significance and type of ploidy-linked differences in their IC_50_ values (Fig. 1A, B, and S3; see also *Materials and methods*). Among the compounds that showed significant ploidy-linked changes in efficacy, a microtubule-stabilizing compound, paclitaxel, had higher efficacy against cells with higher ploidy (hyperploidy-selective; Fig. 1A and S3), consistent with the previous study ^11^. CENP-E inhibitors GSK-923295 and PF-2771 were also remarkably hyperploidy-selective (Fig. 1A and S3). Hyperploidy-selective suppression by CENP-E inhibitors was also observed in another tetraploid HAP1 cell line (Fig. S4A and B). As previously reported ^13^, a Plk1 inhibitor, BI-2536, suppressed tetraploid cells more efficiently than diploids, while its efficacy was equivalent between haploids and diploids. In contrast, an importin-β inhibitor importazole and an Eg5 inhibitor S-trityl-L-cysteine (STLC) had higher efficacy against cells with lower ploidy (hypoploidy-selective; Fig. 1A and S3). Consistent with STLC, another Eg5 inhibitor, monastrol suppressed haploid cells more efficiently than diploids, while its efficacy was equivalent between diploids and tetraploids (Fig. S3). Topoisomerase II inhibitors, daunorubicin, doxorubicin, and etoposide tended to suppress the proliferation of cells with different ploidies with equivalent efficacy. Diverse profiles of ploidy-linked changes in the efficacy of different anti-mitotic compounds indicate that ploidy difference has complex and non-uniform effects on different aspects of molecular regulations of cell division. The ploidy-linked change in the efficacy of CENP-E inhibitors was particularly notable and previously unreported. Therefore, we decided to address further the significance and mechanism of tetraploidy selectivity of CENP-E inhibitors in comparison with paclitaxel, a previously reported tetraploidy-selective compound ^11^.

**Fig. 1:**
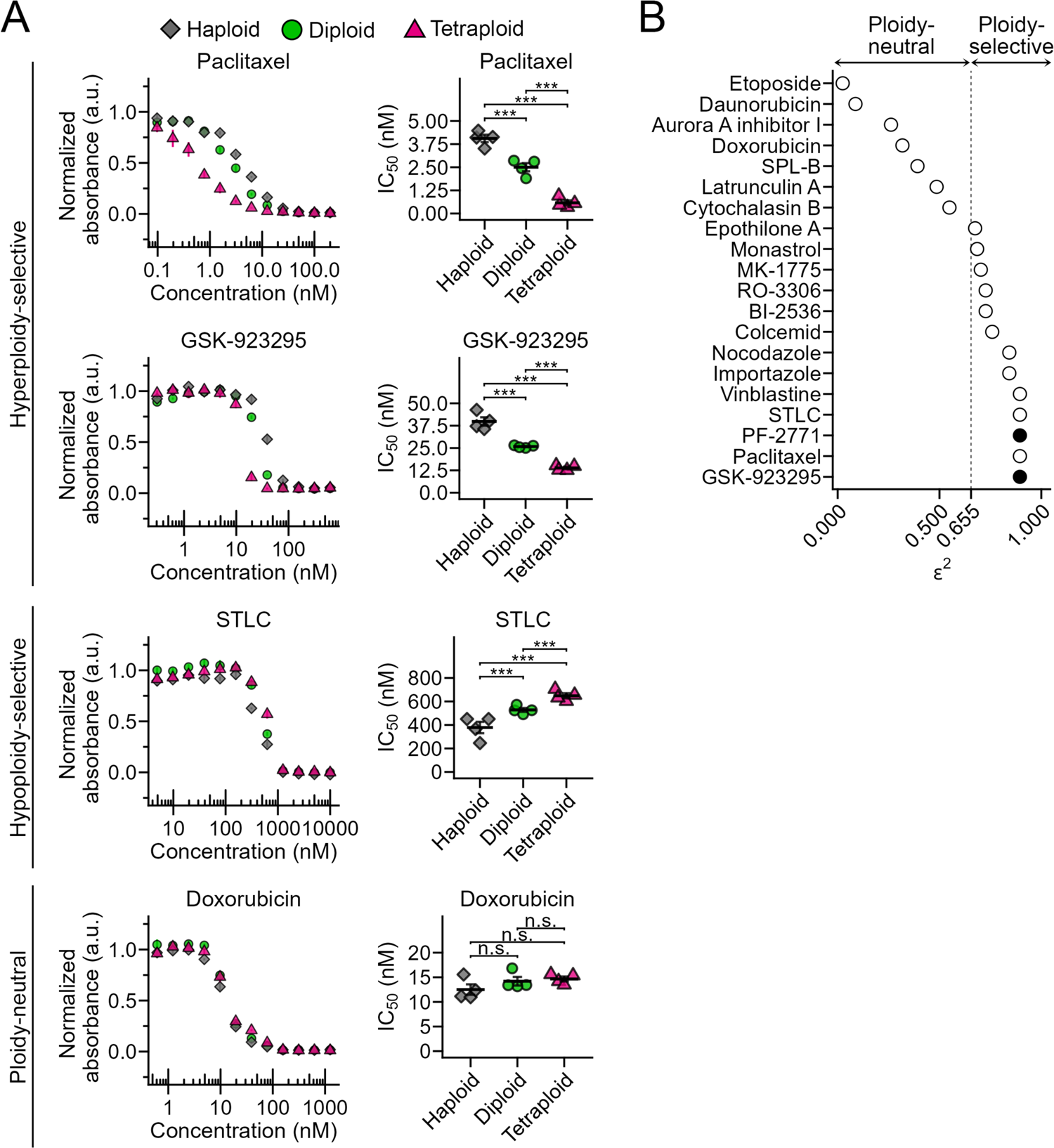
Identification of ploidy-selective anti-mitotic compounds. (**A**) Dose-response curve of normalized absorbance (left) and calculated IC_50_ values (right) in a comparative colorimetric cell proliferation assay using anti-mitotic compounds in haploid, diploid, and tetraploid HAP1 cells. Mean ± standard error (SE) of 4 samples from 2 independent experiments for each condition. Asterisks indicate statistically significant differences in IC_50_ between cells with different ploidies (\*\*\**p* < 0.001, n.s.: not significant, the Steel-Dwass test). See also Fig. S2 and 3 for data of all compounds tested. (**B**) Evaluation of ploidy selectivity of different anti-mitotic compounds based on effect size *ε^2^* of ploidy-linked IC_50_ differences calculated by the Kruskal-Wallis test. The filled circles indicate CENP-E inhibitors.

**Fig. 2:**
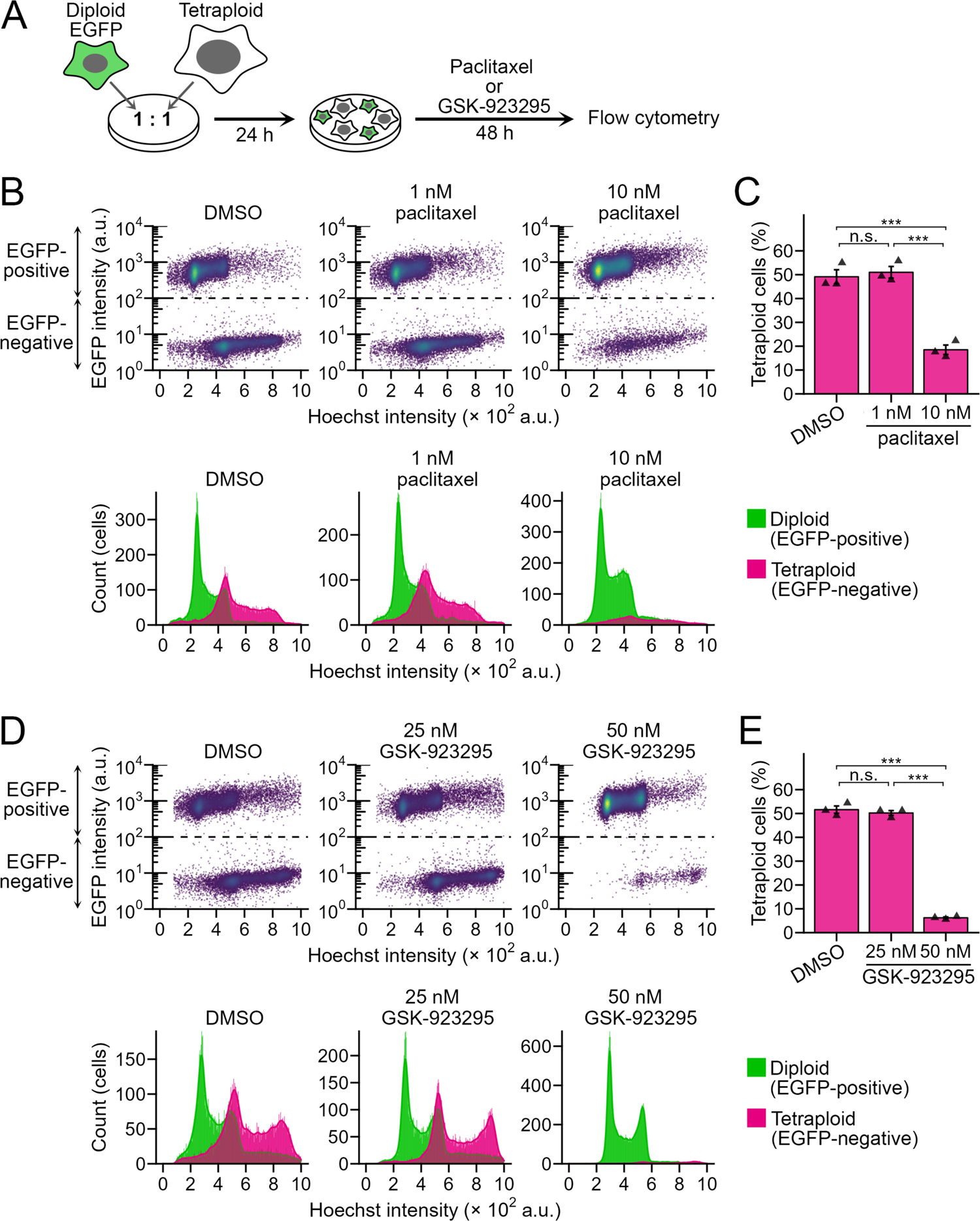
Selective suppression of tetraploid HAP1 cells in diploid-tetraploid co-culture by paclitaxel or GSK-923295. (**A**) Scheme of diploid-tetraploid co-culture experiment. (**B**, **D**) Flow cytometric analyses of diploid and tetraploid cell numbers in their co-culture treated with paclitaxel (B) or GSK-923295 (D) for 48 h. Dot plots of EGFP intensity against the Hoechst signal (corresponding to DNA content) or histograms of the Hoechst signal are shown on top or bottom, respectively. Cell populations originating from diploid or tetraploid cells were distinguished based on EGFP signal intensity and separately displayed in the histograms. (**C**, **E**) The proportion of tetraploid cells in the diploid-tetraploid co-culture. Mean ± SE of 3 independent experiments for each condition. Asterisks indicate statistically significant differences between conditions (\*\**\*p* < 0.001, the Steel-Dwass test).

We next investigated the effect of paclitaxel and GSK-923295 on cell proliferation in 1:1 co-culture of EGFP-labeled diploid and unlabeled tetraploid HAP1 cells (Fig. 2A and S1A). Flow cytometric analysis revealed that DMSO-treated co-culture roughly kept the original diploid-tetraploid ratio after 48-h treatment (Fig. 2B-E), demonstrating that diploid and tetraploid cells proliferated at a similar rate in this condition. On the other hand, 10 nM paclitaxel or 50 nM GSK-923295 significantly reduced tetraploid proportion in the co-culture (tetraploid cells reduced to 19% or 6.2%, respectively; Fig. 2B-E), illustrating the high potential of CENP-E as a target for tetraploidy-selective suppression within heterogeneous cell populations.

### Tetraploidy-linked aggravation of chromosome misalignment, mitotic arrest, and subsequent cohesion fatigue upon CENP-E inhibition

To understand the cause of the tetraploidy-selective growth suppression by CENP-E inhibition, we conducted live imaging of the mitotic progression in co-cultured diploid and tetraploid HAP1 cells. Diploid and tetraploid cells were differentially labeled by stably expressing histone H2B transgene tagged with EGFP and mCherry, respectively (Fig. 3A, B, and S1A). In DMSO-treated co-culture, diploid and tetraploid cells underwent normal cell division with an average mitotic duration of 34 and 30 min (from NEBD to anaphase onset), respectively (Fig. 3C and D). When treated with 50 nM GSK-923295, which caused sharp tetraploidy-selective suppression (Fig. 2E), diploid and tetraploid cells manifested misaligned polar chromosomes at a high frequency in the early mitotic stage (85% and 100% of diploid and tetraploid cells, respectively; Fig. 3B, E, and F). In most GSK-923295-treated diploid cells, these polar chromosomes gradually moved into the metaphase plate, and all chromosomes eventually aligned (Fig. 3B and C). As a result, the majority of diploid cells (87%) entered anaphase and completed cell division despite considerable mitotic delay (with an average mitotic duration of 197 min; Fig. 3D, and G). Compared to diploids, GSK-923295-treated tetraploid cells manifested severer polar chromosome misalignment (Fig. 3E and F). In most cases, these polar chromosomes also gradually moved into the metaphase plate but never completed the alignment (Fig. 3B, S5A, and B). As a result, GSK-923295-treated tetraploid cells spent an extremely long time in mitosis (with an average mitotic duration of 713 min), and 87% of them eventually underwent cohesion fatigue (the catastrophic chromosome scattering) ^24^ (Fig. 3B, C, and H). Cohesion fatigue took place 347 ± 15 min after NEBD (mean ± standard error, n=53 from 2 independent experiments) in GSK-923295-treated tetraploid cells when most GSK-923295-treated diploids had completed congression of initially misaligned chromosomes and entered anaphase (Fig. 3C). Subsequently to cohesion fatigue, GSK-923295-treated tetraploid cells either died during mitosis or exit mitosis without chromosome segregation (mitotic slippage; Fig. 3B and G). A substantial proportion of GSK-923295-treated tetraploid cells (63%) that exit mitosis died during the next cell cycle (Fig. 3I). In contrast, most GSK-923295-treated diploids survived through the next cell cycle despite the delay in the previous mitosis. These results suggest that the ploidy-dependent difference in time duration of mitotic arrest critically affects the fate of CENP-E-inhibited cells: While diploid cells resolve mitotic arrest within the critical time window for chromatid cohesion maintenance in the above CENP-E inhibitory condition, tetraploids go beyond that time window and suffer catastrophic mitotic damages.

**Fig. 3:**
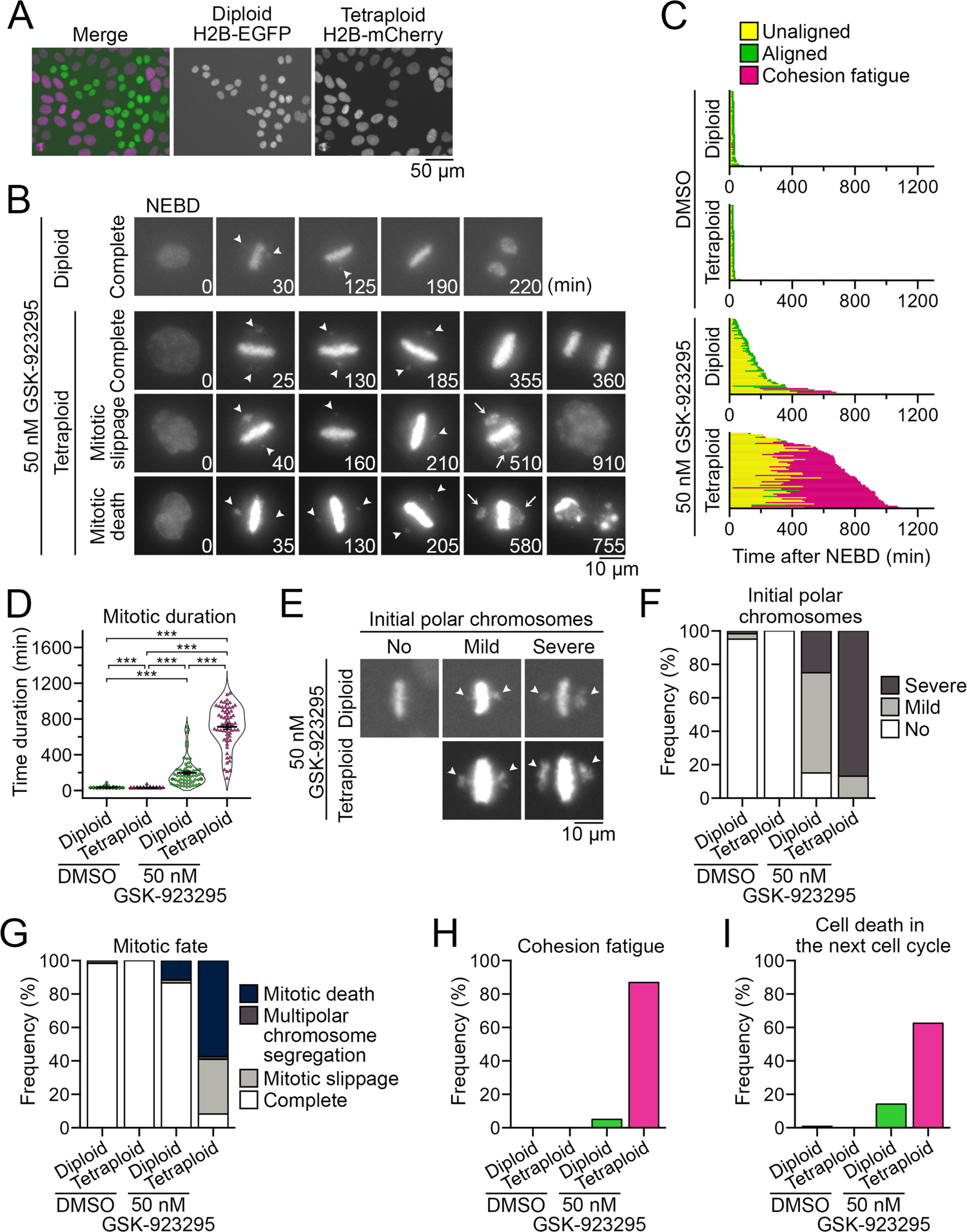
Tetraploidy-linked aggravation of chromosome misalignment and mitotic failure upon GSK-923295 treatment. (**A**) Fluorescence microscopy of co-cultured diploid and tetraploid HAP1 cells expressing histone H2B-EGFP and histone H2B-mCherry, respectively. (**B**) Time-lapse images of the mitotic progression of GSK-923295-treated diploid or tetraploid cells in the co-culture. Arrowheads: misaligned polar chromosomes. Arrows: Gross chromosome scattering caused through cohesion fatigue. (**C**) Analysis of mitotic progression of control and GSK-923295-treated diploid or tetraploid cells in B. Each bar represents a single mitotic event (from NEBD to anaphase onset or mitotic exit) in a dividing cell. At least 60 cells from 2 independent experiments were analyzed for each condition. (**D**) Mitotic duration (from NEBD to anaphase onset or mitotic exit) in control and GSK-923295-treated diploid or tetraploid cells in B. Mean ± SE of at least 60 cells from 2 independent experiments for each condition. Asterisks indicate statistically significant differences between conditions (\*\**\*p* < 0.001, the DSCF test). (**E**) Different degrees of polar chromosome misalignment appeared upon the formation of the metaphase plates (initial polar chromosomes; arrowheads) in GSK-923295-treated diploid or tetraploid cells. (**F-I**) Frequency of different degrees of initial polar chromosome misalignment (F), mitotic fates (G), cohesion fatigue event (H), or cell death in the subsequent cell cycle (I) in control and GSK-923295-treated diploid or tetraploid cells in B. At least 60 cells (F-H) or 32 cells (I) from 2 independent experiments were analyzed for each condition.

A recent study revealed that tetraploid cells were particularly defective in retention of pre-aligned metaphase chromosomes upon inhibition of a mitotic kinesin Kif18A, highlighting the unstable nature of metaphase plate in tetraploid cells ^10^. This prompted us to test the effect of CENP-E inhibition on the retention of pre-aligned chromosomes in diploid and tetraploid cells. For this, we used a previously developed photo-switchable CENP-E inhibitor (PCEI-HU), which reversibly converts to non-inhibitory *cis* or inhibitory *trans* isomer by irradiating UV or visible light, respectively ^25^ (Fig. S6A). Diploid and tetraploid cells were treated with the inhibitor at the photo-stationary state (PSS) enriched in the non-inhibitory *cis* isomer along with MG132 and SiR-DNA (for blocking anaphase onset and staining mitotic chromosomes, respectively) for 2 h. Then mitotic chromosomes were live imaged (see *Materials and methods*). During the live imaging, the inhibitor was switched to the PSS enriched in the inhibitory *trans* isomer by irradiating 505 nm light. The photo-switching of the inhibitor in prometaphase cells that still possessed unaligned chromosomes resulted in misaligned polar chromosomes, demonstrating that the inhibitor was indeed switched to the inhibitory state after the photo-irradiation (Fig. S6B). In contrast, photo-switching of the inhibitor in metaphase cells in which all chromosomes aligned at the equatorial plate, *de novo* misalignment of the pre-aligned chromosomes was seldom observed either in diploids or tetraploids (Fig. S6C-F). This result indicates that aggravation of initially formed misaligned chromosomes rather than failure to maintain pre-aligned chromosomes is likely to cause extremely prolonged mitosis in CENP-E-inhibited tetraploid cells.

### Tetraploidy-linked aggravation of spindle multipolarization and subsequent cell death by paclitaxel treatment

Previous studies revealed that paclitaxel’s effects on mitotic control are pleiotropic and concentration-dependent ^26–29^, and cellular processes of the tetraploidy-selective suppression by paclitaxel remained unclear. To specify the paclitaxel-induced mitotic defects aggravated by tetraploidy and gain insight into the cellular basis of tetraploidy-selective growth suppression, we compared the effect of paclitaxel on the mitotic progression of co-cultured diploid and tetraploid cells (Fig. 4A). In the presence of 10 nM paclitaxel, which caused tetraploidy-selective suppression in co-culture (Fig. 2C), mitotic progression was significantly delayed in tetraploid cells (with an average mitotic duration of 490 min or 32 min in paclitaxel- or DMSO-treated tetraploid cells, respectively; Fig. 4B and C). The paclitaxel-induced mitotic delay was milder in diploid cells (with an average mitotic duration of 91 min or 37 min in paclitaxel- or DMSO-treated diploid cells, respectively; Fig. 4C). Importantly, most paclitaxel-treated tetraploid cells (97%) manifested Y-shaped abnormal metaphase plates, frequently followed by multipolar chromosome segregation, mitotic death or mitotic slippage (Fig. 4A, B, and D). The majority of paclitaxel-treated tetraploid cells that exited mitosis died during the next cell cycle (Fig. 4E). These mitotic defects were much less frequent in paclitaxel-treated diploids, and most of them underwent normal bipolar chromosome segregation and survived through the next cell cycle (Fig. 4D and E). These results suggest that the tetraploidy-linked aggravation of multipolar division is a primary cause of tetraploidy-selective growth suppression by paclitaxel.

**Fig. 4:**
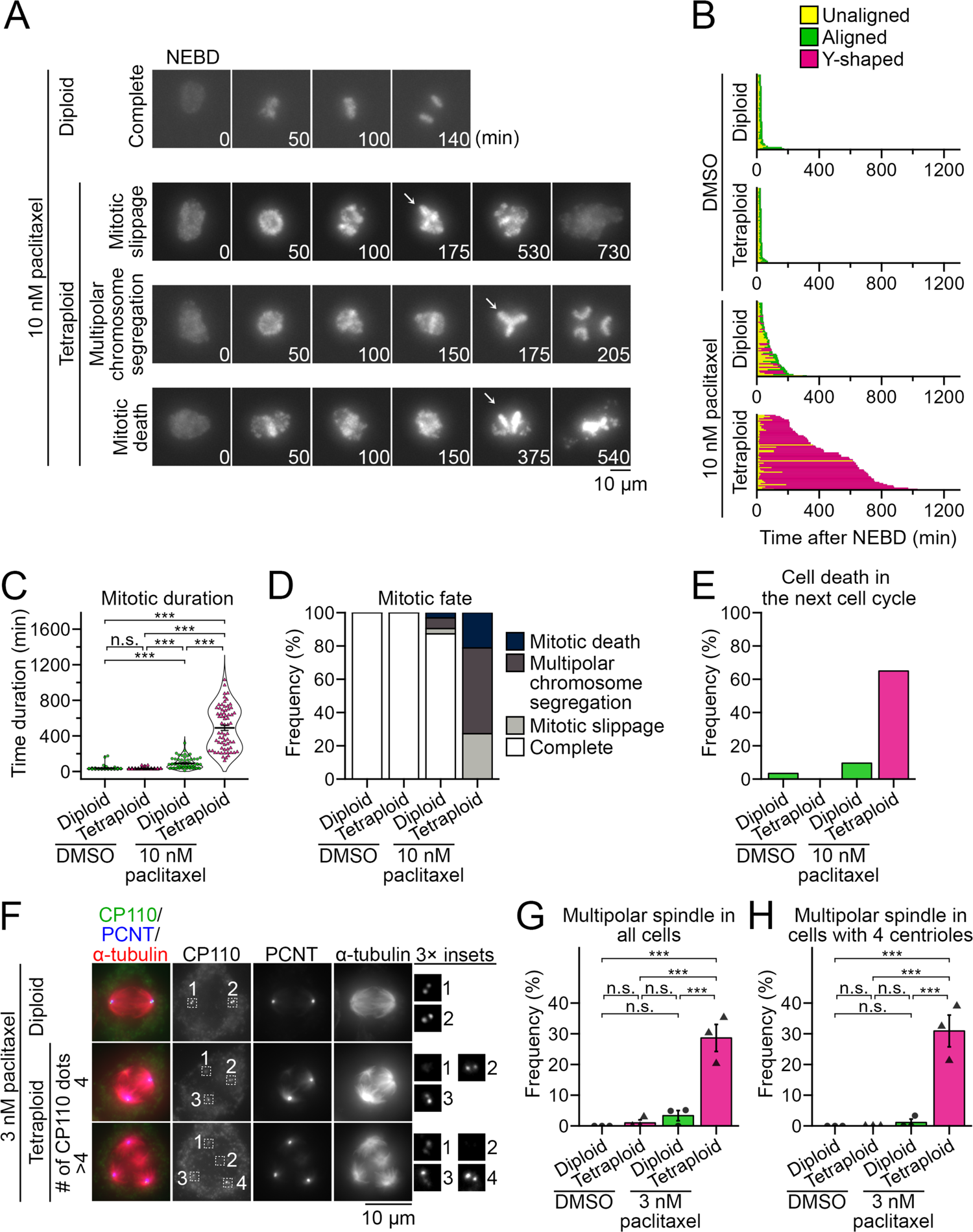
Tetraploidy-linked aggravation of multipolar spindle formation upon paclitaxel treatment. (**A**) Time-lapse images of the mitotic progression in paclitaxel-treated diploid H2B-EGFP and tetraploid H2B-mCherry HAP1 co-culture. Arrows: Y-shaped chromosome arrangement. (**B**) Analysis of mitotic progression of control and paclitaxel-treated diploid or tetraploid cells in A. Each bar represents a single mitotic event (from NEBD to anaphase onset or mitotic exit) in a dividing cell. At least 59 cells from 2 independent experiments were analyzed for each condition. (**C**) Mitotic duration (from NEBD to anaphase onset or mitotic exit) in control and paclitaxel-treated diploid or tetraploid cells in A. Mean ± SE of at least 59 cells from 2 independent experiments for each condition. Asterisks indicate statistically significant differences between conditions (\*\**\*p* < 0.001, the DSCF test). (**D**, **E**) Frequency of mitotic fates (D), or cell death in the subsequent cell cycle (E) in control and paclitaxel-treated diploid or tetraploid cells in A. At least 59 or 97 cells from 2 independent experiments were analyzed for each condition in D or E, respectively. (**F**) Immunofluorescence microscopy of CP110, PCNT, and α-tubulin in 3 nM paclitaxel-treated diploid or tetraploid cells. (**G**, **H**) Frequency of multipolar spindle in control and paclitaxel-treated diploid or tetraploid cells in F. Data obtained from all cells or only cells with 4 centrioles were shown in G or H, respectively. Mean ± SE of 3 independent experiments. At least 92 or 90 cells were analyzed for each condition in G or H, respectively. Asterisks indicate statistically significant differences between conditions (\*\**\*p* < 0.001, the Steel-Dwass test).

Multipolar chromosome segregation accompanying the formation of a “Y-shaped” metaphase plate suggests spindle multipolarization during pre-anaphase in the paclitaxel-treated tetraploid cells. To test this possibility, we conducted immunostaining against α-tubulin, pericentrin, and CP110 (makers of microtubules, pericentriolar material, and the centrioles, respectively) in DMSO- or 3 nM paclitaxel-treated diploid or tetraploid cells (Fig. 4F-H). Previously, we found that tetraploid cells suffered chronic centriole overduplication ^23^. Therefore, to distinguish the direct influence of tetraploidy on spindle polarity upon paclitaxel treatment from indirect one through the formation of extra centrosomes, we sorted cells based on the centriole number per cell in the spindle polarity analysis (Fig. 4H). Paclitaxel-treated tetraploid cells possessed multipolar spindle at a significantly higher frequency than DMSO-treated control or paclitaxel-treated diploid cells, either when all cells or only the cells possessing 4 centrioles were counted in the quantification (Fig. 4G and H). This result suggests that tetraploidy per se, rather than the presence of extra centrosomes, promotes the spindle multipolarization upon low concentration paclitaxel treatment, making tetraploid cells more prone to lethal chromosome loss.

### CENP-E inhibitor shows selectivity toward a broader spectrum of tetraploid cell lines than paclitaxel

The above results indicate that CENP-E inhibitor and paclitaxel selectively suppress tetraploid cell proliferation through different mechanisms, prompting us to compare their effects on tetraploid cells with different cellular backgrounds. For this, we investigated the effect of paclitaxel and CENP-E inhibitors on the viability of another near-diploid human cell line, HCT116, and 16 isogenic tetraploid lines (Fig. 5A and B, S1B and S7). We found variation in the efficacy of paclitaxel among different tetraploid HCT116 cell lines: While paclitaxel suppressed 12 tetraploid cell lines significantly more efficiently than diploid, its IC_50_ values dispersed among these lines (Fig. 5A). In the remaining 4 tetraploid cell lines, the efficacy of paclitaxel did not significantly differ from that in diploids. This result indicates the limited generality of the tetraploidy selectivity of paclitaxel. On the other hand, GSK-923295 had significantly higher efficacy against all 16 tetraploid HCT116 lines than diploids with IC_50_ values comparable among these tetraploid lines (Fig. 5B), highlighting consistent selectivity of CENP-E inhibition towards tetraploid cells in different backgrounds.

**Fig. 5:**
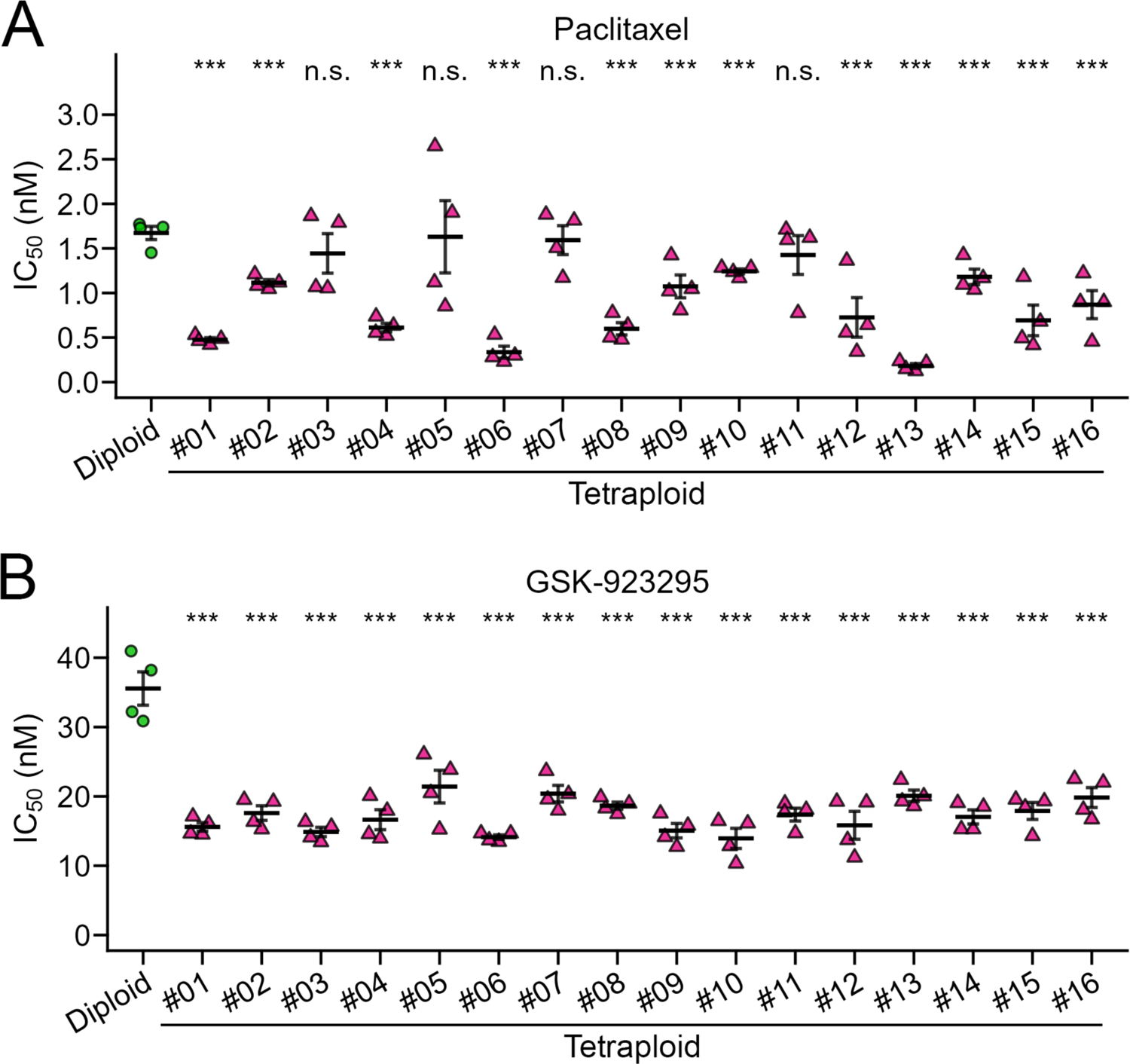
Comparison of efficacy of paclitaxel or GSK-923295 in different tetraploid HCT116 lines. (**A**, **B**) IC_50_ values in a comparative colorimetric cell proliferation assay using paclitaxel (A) and GSK-923295 (B) in diploid or 16 different tetraploid HCT116 cell lines. Mean ± SE of 4 samples from 2 independent experiments for each condition. Asterisks indicate statistically significant differences in IC_50_ between the control diploid line and each tetraploid line (\*\*\**p* < 0.001, the Steel test). See also Fig. S7 for the dose-response curve of normalized absorbance used for calculating IC_50_.

## Discussion

Ploidy alteration causes pleiotropic changes in cell structures and contents, including chromosome number, cell volume, or whole-protein amount, having a profound quantitative effect on mitotic machinery ^13, 23, 30^. However, the effects of ploidy alteration on the molecular function of mitotic regulators remain largely unknown. This study revealed that ploidy alteration changes cellular sensitivity to different anti-mitotic compounds in a complex and non-uniform manner. Among these compounds, CENP-E inhibitors showed remarkable and consistent hyperploidy selectivity in mitotic perturbation and cell proliferation suppression through a different mechanism than a previously reported hyperploidy-selective compound, paclitaxel. CENP-E inhibition manifested superior consistency to paclitaxel in the tetraploidy selectivity across cell lines, suggesting its potential utility in tetraploidy-specific suppression in a broad spectrum of cellular backgrounds.

Our results indicate that the tetraploidy-linked aggravation of mitotic failure is the leading cause of the sharp tetraploidy selectivity of low-dose CENP-E inhibition (Fig. 3). Based on our live imaging, we propose that the tetraploidy-linked aggravation of mitotic failure upon CENP-E inhibition stems from the combination of i) the tetraploidy-linked increase in chromosome misalignment and ii) cohesion fatigue frequently occurring in the time gap between mitotic exit in diploids and tetraploids. To explain point i) above, we speculate that the doubled chromosome number is the direct cause of the aggravation of chromosome misalignment in CENP-E-inhibited tetraploid cells. A previous study reported that CENP-E mediates the congression of only a subset of chromosomes located in peripheral areas within the nucleus upon the mitotic entry ^21^. The doubled chromosome number with the enlarged nucleus in tetraploid cells would increase such peripheral chromosomes vulnerable to CENP-E inhibition. Because of the increased polar chromosomes upon CENP-E inhibition, tetraploid cells spent significantly longer time than diploids to solve chromosome misalignment. This differential effect of CENP-E inhibition results in a notable time gap between mitotic exit in diploid and tetraploid cells. To explain point ii) above, cohesion fatigue (premature breakage of sister chromatid cohesion) occurs when mitotic progression is blocked with continuous tension applied at kinetochores of sister chromatids ^31, 32^. Inhibition of CENP-E motor activity satisfies the criteria for inducing cohesion fatigue with its characteristic effects on mitotic regulations: It blocks the congression of a small proportion of chromatids to block mitotic progression by activating SAC (note that upon inhibition of CENP-E activity, CENP-E protein remains at the kinetochores, supporting the recruitment of SAC activation factors) ^18, 33^, while leaving the majority of chromatids aligned at metaphase plate under the tension force exerted by an intact bipolar spindle ^17, 34^. CENP-E-inhibited cells typically undergo cohesion fatigue after >200-min mitotic arrest (Fig. 3C). By that time, most diploid cells resolve chromosome misalignment and exit mitosis. In contrast, most tetraploid cells remain at mitosis with unsolved chromosome misalignment and undergo irreversible mitotic catastrophe at optimum inhibitor concentration. Based on this model of tetraploidy-selective suppression, it would be intriguing to address potential ploidy selectivity of different interventions that satisfy the criteria described above: The interventions that differentially modulate mitotic progression among different ploidies while facilitating the occurrence of cohesion fatigue.

We also found that tetraploid cells are more prone to spindle multipolarization than diploid cells upon paclitaxel treatment. Notably, the paclitaxel concentration most effective for tetraploid-selective suppression was within the clinically relevant range of the drug concentration ^28^. The cause of the tetraploidy-linked increase in spindle multipolarization remains unknown. Interestingly, a recent study reported that polyploid drosophila embryonic cells were more prone to spindle multipolarization because of the increased steric hindrance of the polyploid amount of chromosomes that precludes the supernumerary centrosomes from clustering into bipolar spindle poles ^35^. Spindle multipolarization frequently took place even in the tetraploid cells with the normal centrosome number (Fig. 4H), indicating that the tetraploidy-linked aggravation of spindle multipolarity upon paclitaxel treatment occurs by a different mechanism than the one depending on supernumerary centrosomes. We speculate that drastic changes in quantitative features of the mitotic spindle may make tetraploid cells more prone to multipolarize upon paclitaxel treatment. Future studies would provide further insight into the molecular basis of the tetraploidy selectivity of paclitaxel and the factors that limit the generality of tetraploidy selectivity among different cellular backgrounds.

A recent study revealed the possibility of selective tetraploid cell suppression by inhibiting Kif18A, whose requirement for maintaining proper alignment of metaphase chromosomes increases in tetraploid cells ^10^. Our study revealed that CENP-E inhibition and paclitaxel selectively suppressed tetraploid cell proliferation through different principles from one another and the previous study. These findings imply that quantitative changes in multifaceted aspects of the mitotic regulatory mechanism upon the whole-genome duplication make tetraploid cells more susceptible to various mitotic perturbations. Moreover, our results demonstrate that different tetraploidy-selective interventions cover a different spectrum of tetraploid cellular backgrounds. Taking the high heterogeneity of tetraploid cells into account ^6^, increasing the choice of drug targets and establishing effective combinations for tetraploid-selective suppression would benefit cancer therapeutics.

## Materials and methods

### Cell culture and flow cytometry

Haploid HAP1 cells ^36^ and their isogenic diploid and tetraploid lines ^23^ were cultured in Iscove’s Modified Dulbecco’s Medium (IMDM; Wako Pure Chemical Industries, Osaka, Japan) supplemented with 10% fetal bovine serum (FBS) and 1× antibiotic-antimycotic solution (AA; Sigma-Aldrich). Haploid cells were maintained by size-based cell sorting as previously described ^23^. HCT116 cells were provided by Riken Cell Bank (RCB2979) and cultured in McCoy’s 5A or Dulbecco’s Modified Eagle Medium (Wako) supplemented with 10% FBS and 1× AA. For establishing tetraploid HCT116 cell lines, diploid cells were treated with 40 ng/mL nocodazole for 4 h, washed 3 times with cell culture medium, shaken off, and treated with 5 μg/mL cytochalasin B for 4 h. Then, cells were washed 3 times with cell culture medium and diluted in 10-cm dishes. After 8-10 d, colonies containing cells that were uniform in size and larger than diploids were clonally expanded and checked for DNA content to select near-tetraploid clones. For DNA content analyses, 2 × 10^6^ cells were stained with 10 μg/ml Hoechst 33342 (Dojindo) for 15 min at 37°C, and DNA content was analyzed using a JSAN desktop cell sorter (Bay bioscience).

### Inhibitors

Inhibitors were purchased from the distributors as follows. Aurora A inhibitor I, BI-2536, epothilone A, and MK-1775: AdooQ BioScience. SPL-B: Axon Medchem. Latrunculin A: Focus Biomolecules. PF-2771: MedChemExpress. GSK-923295: Selleck Chemicals. Importazole, RO-3306, and S-trityl-L-cysteine (STLC): Sigma-Aldrich. Colcemid (KaryoMAX Colcemid): Thermo Fisher Scientific. Etoposide: Calbiochem. Vinblastine: LKT Laboratories. Monastrol: Tocris Bioscience. Cytochalasin B, daunorubicin, doxorubicin, nocodazole, and paclitaxel: Wako.

### Colorimetric cell proliferation assay

For cell viability assay, haploid, diploid, or tetraploid HAP1 cells were seeded on 96-well plates at 2250, 1125, or 562.5 cells/well, respectively. Diploid or tetraploid HCT116 cells were seeded at 1350 or 675 cells/well, respectively. After 24 h, cells were treated with different concentrations of anti-mitotic compounds. Forty-four (HAP1 cells) or 68 h (HCT116 cells) after the addition of the compounds, 5% Cell Counting Kit-8 (Dojindo) was added to the culture, incubated for 4 h, and absorbance at 450 nm was measured using the Sunrise plate reader (Tecan). IC50 was calculated by curve fitting of normalized dose-response data using nonlinear regression:

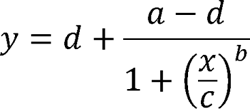

where *y* is the normalized absorbance, *x* is drug concentration, *a* or *d* is the absorbance at zero or infinite drug concentration, respectively, and *b* or *C* is the slope factor or the inflection point, respectively.

### Mixed culture experiment

For flow cytometry analysis, EGFP-labeled diploid and non-labeled tetraploid HAP1 cell suspension (1.5 × 10^4^ cells/ml each) were mixed in a 1:1 ratio, 1.8 ml seeded on 6-well plates coated with collagen type I (Corning). After 24 h, paclitaxel or GSK-923295 was treated in the co-culture. Forty-eight h after the addition of the compounds, cells were trypsinized, suspended in DPBS, stained with 10 μg/ml Hoechst 33342, and analyzed by flow cytometry. The two mixed cell populations were separately counted based on the EGFP fluorescence signal.

For live imaging, diploid and tetraploid cells stably expressing histone H2B transgene tagged with EGFP and mCherry, respectively, were mixed in a 1:1 ratio (1.35 × 10^4^ cells/ml each), 0.2 ml seeded on collagen-coated 8-well imaging chamber. After 24 h, paclitaxel or GSK-923295 was treated in the co-culture, and live imaging was subsequently conducted for 48 h. The first mitotic events after the drug treatment were analyzed.

### Immunofluorescence staining

Cells were fixed with 100% methanol at −20°C for 10 min, treated with BSA blocking buffer (150 mM NaCl, 10 mM Tris-HCl pH 7.5, 5% BSA, and 0.1% Tween 20) for 30 min at 25°C, incubated with rat monoclonal anti-α-tubulin (YOL1/34, EMD Millipore; 1:1000), mouse monoclonal anti-PCNT (ab28144, Abcam; 1:1000), rabbit polyclonal anti-CP110 (A301-343A, Bethyl Laboratories; 1:1000) overnight at 4°C, and with fluorescence (Alexa Fluor 488, 568, 647)-conjugated secondaries (Jackson ImmunoResearch Laboratories or Abcam; 1:1000) overnight at 4°C at indicated dilutions. Following each treatment, cells were washed 3 times with phosphate-buffered saline.

### Microscopy

For fixed cell imaging, cells were observed under a TE2000 microscope (Nikon) equipped with a ×100 1.4 NA Plan-Apochromatic, a CSU-X1 confocal unit (Yokogawa), and an iXon3 electron multiplier-charge coupled device (EMCCD) camera (Andor) or ORCA-ER CCD camera (Hamamatsu Photonics). Live cell imaging was conducted at 37°C with 5% CO_2_ using a Ti-2 microscope (Nikon) equipped with ×20 0.75 NA Plan-Apochromatic, and Zyla4.2 sCMOS camera (Andor). For live imaging, cells were cultured in phenol red-free IMDM (Thermo Fisher Scientific) supplemented with 10% FBS and 1× AA. Image acquisition was controlled by µManager (Open Imaging).

### Photo-switching CENP-E inhibition experiment

One mM stock solution of PCEI-HU, a photo-switchable CENP-E inhibitor, in dimethyl sulfoxide was diluted at 1:2000 in IMDM in a microtube, then irradiated with 365 nm LED light (Asahi Spectra, 416 mW/cm^2^ at 100%, irradiated from 5 cm above the sample for 60 s) to reach a photostationary state (PSS) enriched in non-inhibitory *cis* isomer, and immediately treated in diploid or tetraploid HAP1 cells at the final concentration of 0.5 µM. At the same time, cells were co-treated with 10 µM MG132 (Peptide Institute; for blocking anaphase onset) and 100 nM SiR-DNA (Cytoskeleton inc.; for visualizing mitotic chromosomes). After 2-h incubation in the dark, we started far-red fluorescence live imaging of SiR-DNA-stained mitotic chromosomes. Note that observing light for live imaging does not affect the photoisomerization of PCEI-HU ^25^. At 15 min after the initiation of live imaging, PCEI-HU-treated cells were irradiated with 505 nm LED light (Asahi Spectra, 141 mW/cm^2^ at 100%, irradiated from 3.2 cm above the sample for 35 s) to make the compound reach a PSS enriched in inhibitory *trans* isomer of PCEI-HU. We then traced the motion of mitotic chromosomes pre-aligned at the metaphase plate at the time of 505-nm light irradiation.

### Statistical analysis

All data subjected to statistical analyses in this study were abnormally distributed in the Shapiro-Wilk test. For comparing two data groups not assumed to have equal variances, we used the Brunner-Munzel test. For comparing more than two groups of data with equal or unequal sample sizes, we used the Steel-Dwass test or the Dwass-Steel-Critchlow-Fligner (DSCF) test, respectively. In the case of comparing a common diploid control with each of multiple tetraploid samples (Fig. 5), we used the Steel test. Multiple group analyses of drug IC_50_ differences among haploid, diploid, and tetraploid cells (Fig. 1B) were conducted using the Kruskal-Wallis test with post-hoc Steel-Dwass test. Statistical significance was set at *p* < 0.05 for all analyses. The compounds with the effect size of Kruskal-Wallis test ε^2^ > 0.655 were defined as “significantly ploidy-selective” in Fig. 1B (Albers and Lakens, 2018). All statistical analyses were conducted with R software (4.2.1) using brunnermunzel, minpack.lm, PMCMRplus, rcompanion, Rmisc, nparcomp, rstatix, and stats packages.

## Acknowledgment

We are grateful to Sarada Bulchand for commenting on the manuscript, Gohta Goshima for sharing reagents, Kenichi Shimada for technical instruction, and the Global Facility Center at Hokkaido University for the use of flow cytometer. This work was supported by the Sasakawa Scientific Research Grant and Hokkaido University-Hitachi Joint Cooperative Support Program for Education and Research to K. Yo., JSPS KAKENHI (Grant Numbers JP19J12210, and JP21K20737 to K.Ya., and JP19KK0181, JP19H05413, JP19H03219, JPJSBP120193801, and JP21K19244 to R.U.), the Princess Takamatsu Cancer Research Fund, the Kato Memorial Bioscience Foundation, the Orange Foundation, the Smoking Research Foundation, Daiichi Sankyo Foundation of Life Science, and the Nakatani Foundation to R.U. The authors declare no competing financial interests.

## Author Contributions

Conceptualization, K.Yo., M.S., and R.U.; Methodology, K.Yo., A.M., M.S., K.Ya., K.M., and R.U.; Investigation, K.Yo., A.M., M.S., T.Y., and R.U.; Formal Analysis, K.Yo., A.M., and M.S.; Resources, K.Yo., S.I., E.K., T.K., K.M., N.T., R.S., M.M., and R.U.; Writing – Original Draft, K.Yo., and R.U.; Writing – Review & Editing, K.Yo, A.M., and R.U.; Funding Acquisition, K.Yo., K.Ya., M.M., and R.U.

---

**Fig. S1:**
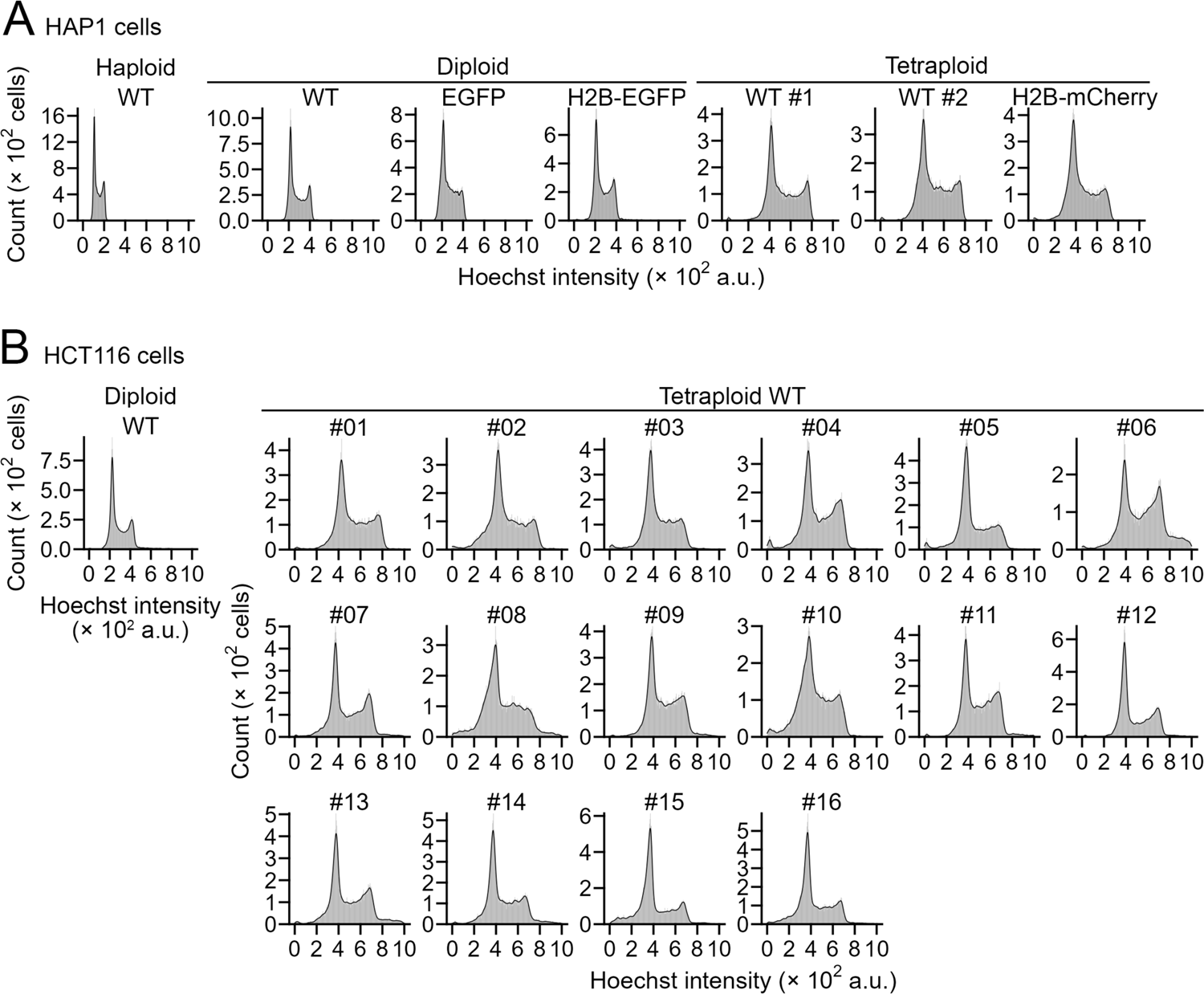
DNA content analyses of cell lines used in this study. (**A**, **B**) Histograms of Hoechst signal in haploid HAP1 cells and their isogenic diploid and tetraploid lines (A), or diploid HCT116 cells and their isogenic tetraploid lines (B). Representative data from 2 independent experiments.

**Fig. S2:**
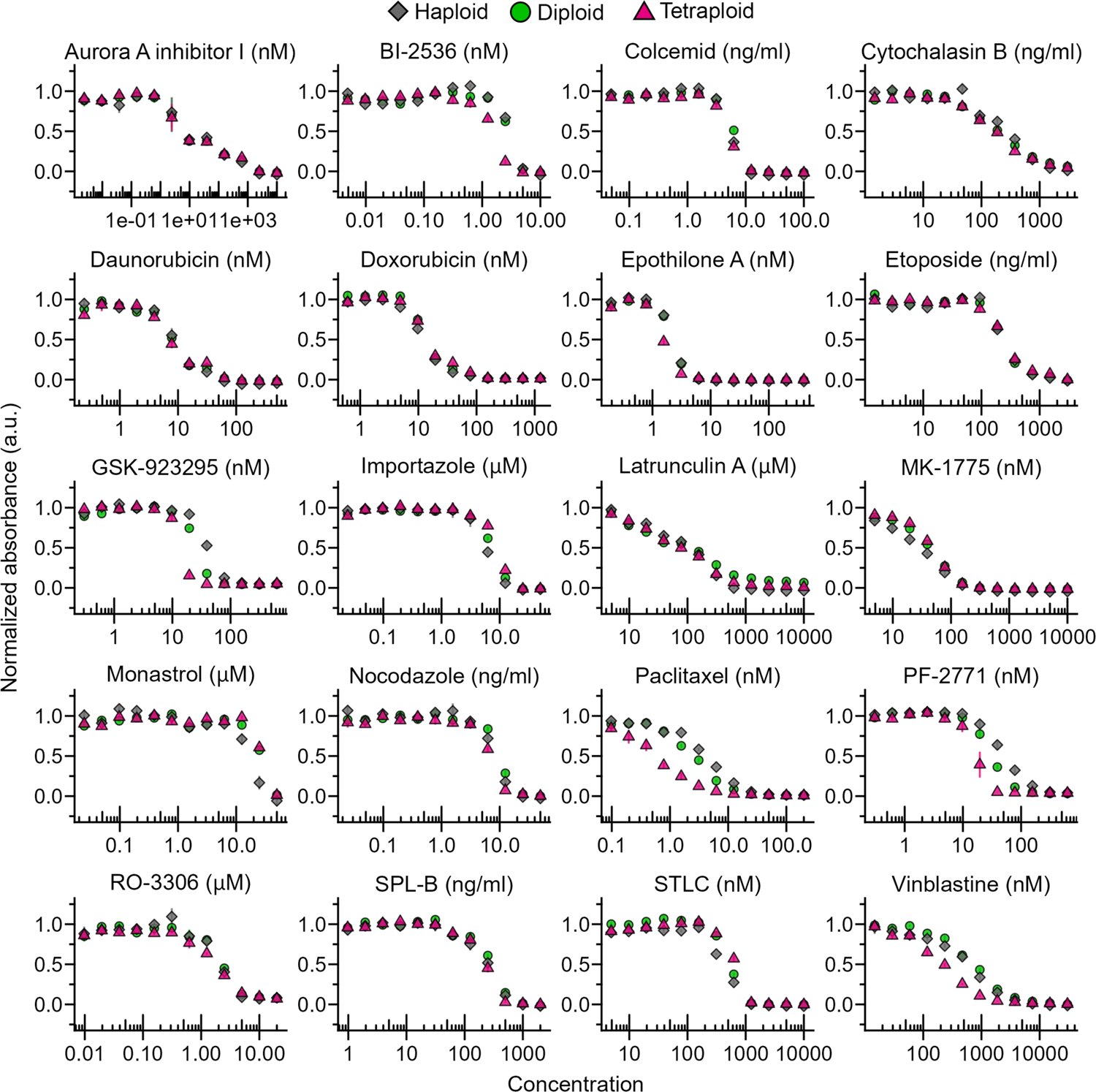
Proliferation of haploid, diploid, or tetraploid HAP1 cells treated with different concentrations of anti-mitotic compounds Dose-response curve of normalized absorbance in a comparative colorimetric cell proliferation assay using different anti-mitotic compounds in haploid, diploid, and tetraploid HAP1 cells. Mean ± SE of 4 samples from 2 independent experiments for each condition. Unit of inhibitor concentration is shown on the top of each graph. The identical data on paclitaxel, GSK-923295, STLC, and doxorubicin were also shown in Fig. 1A.

**Fig. S3:**
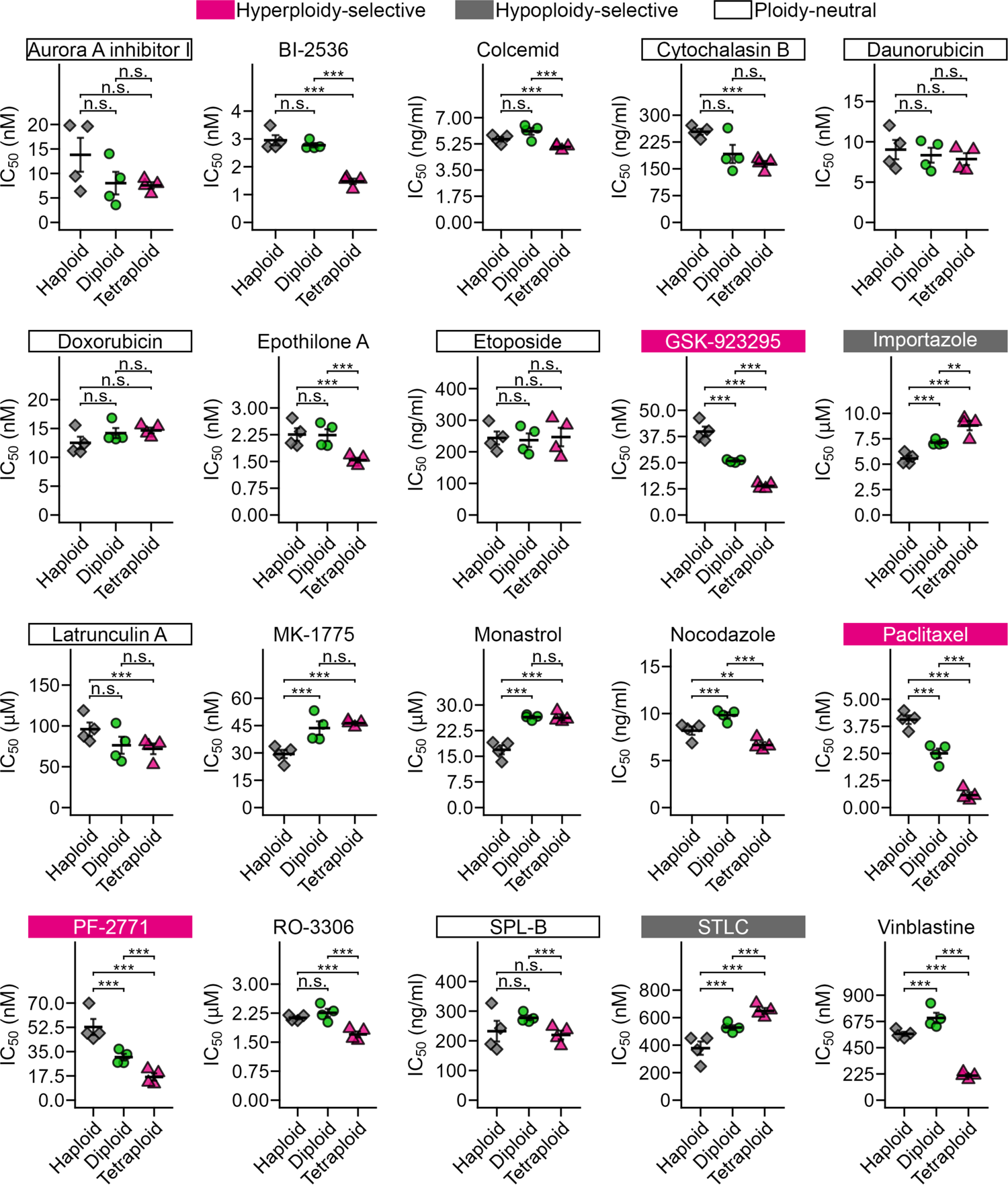
Ploidy-dependent changes in anti-mitotic compound efficacy. IC_50_ values of anti-mitotic compounds in haploid, diploid, and tetraploid HAP1 cells (calculated from the dose-response curves in Fig. S2). Mean ± SE of 4 samples from 2 independent experiments for each condition. Asterisks indicate statistically significant differences in IC_50_ between cells with different ploidies (\*\**p* < 0.01, \*\*\**p* < 0.001, the Steel-Dwass test). The identical data on paclitaxel, GSK-923295, STLC, and doxorubicin were also shown in Fig. 1A. Inhibitors that have significant ploidy-dependent differences in their efficacy (effect size *ε^2^* > 0.655 in the Kruskal-Wallis test; see also Fig. 1B) with positive or negative linear correlations are categorized as “hyperploidy- or hypoploidy-selective,” respectively. Inhibitors with no significant ploidy-dependent differences in efficacy in the Kruskal-Wallis test are categorized as “ploidy-neutral.”

**Fig. S4:**
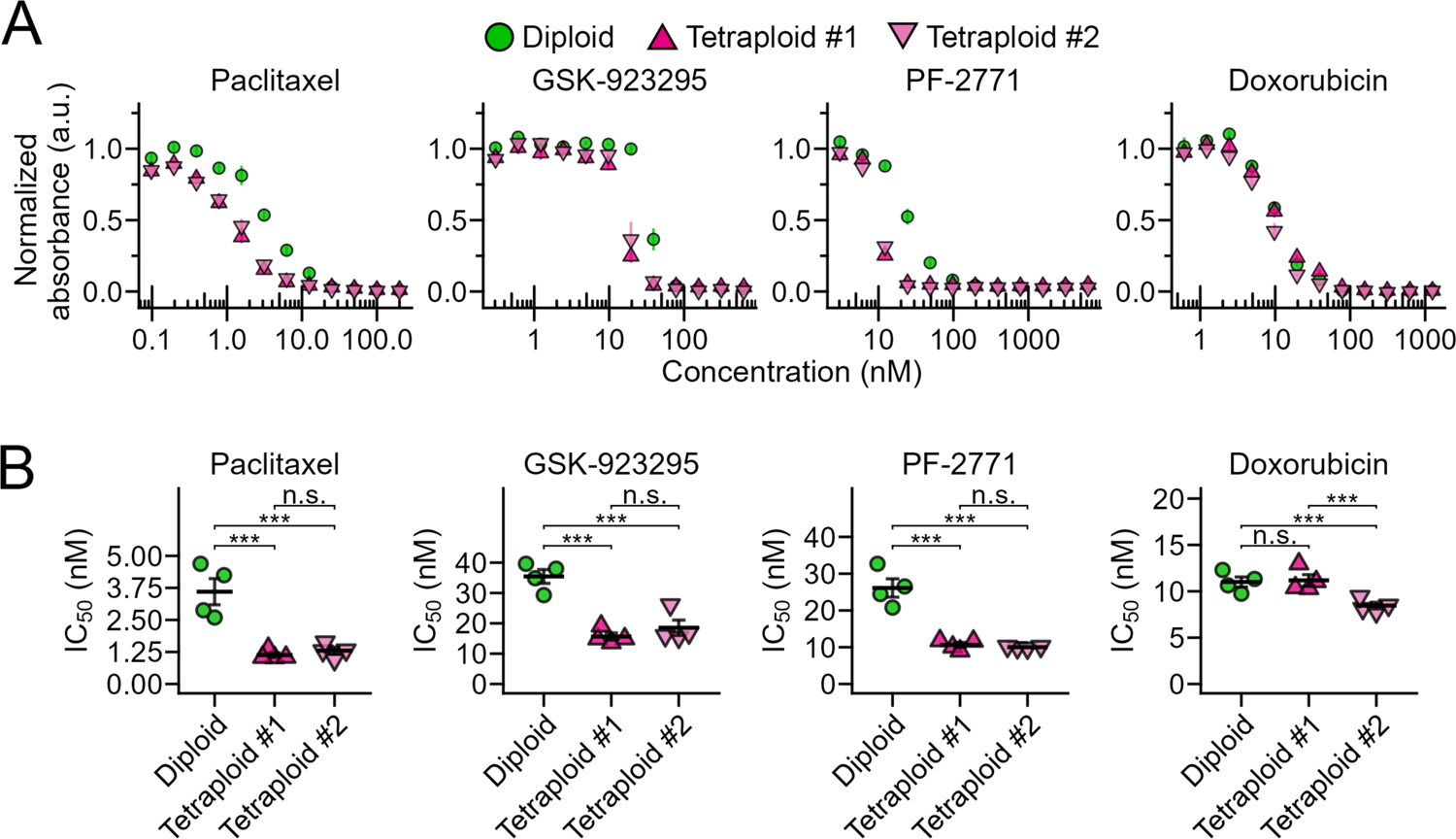
Selective anti-proliferative effect of paclitaxel and CENP-E inhibitors toward 2 independent HAP1 tetraploid cell lines. (**A**, **B**) Dose-response curve of normalized absorbance (A) and calculated drug IC_50_ values (B) in a comparative colorimetric cell proliferation assay using paclitaxel, CENP-E inhibitors, or doxorubicin in diploid and 2 different tetraploid HAP1 cell lines. Mean ± SE of 4 samples from 2 independent experiments for each condition. Asterisks indicate statistically significant differences in IC_50_ between cells with different ploidies (\*\*\**p* < 0.001, the Steel-Dwass test).

**Fig. S5:**
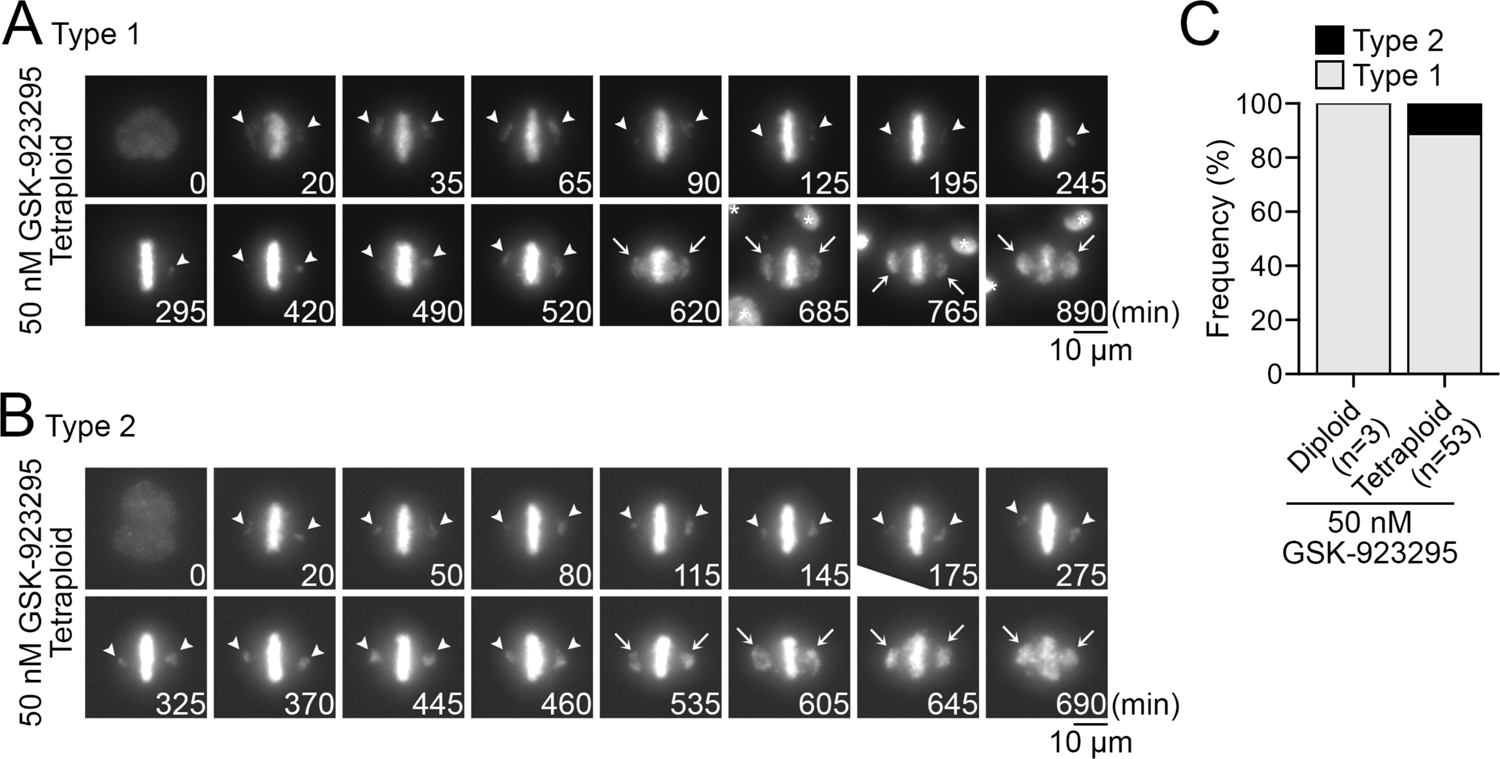
Gradual re-alignment of misaligned polar chromosomes in GSK-923295-treated cells. (**A**, **B**) GSK-923295-treated tetraploid cells whose polar chromosomes gradually moved into the metaphase plate (A; type 1) or did not undergo re-alignment (B; type 2). Arrowheads: misaligned polar chromosomes. Arrows: Gross chromosome scattering caused through cohesion fatigue. (**C**) Frequency of different types of misaligned chromosome movement before cohesion fatigue in GSK-923295-treated diploid or tetraploid cells. Cells that underwent cohesion fatigue were analyzed from 2 independent experiments.

**Fig. S6:**
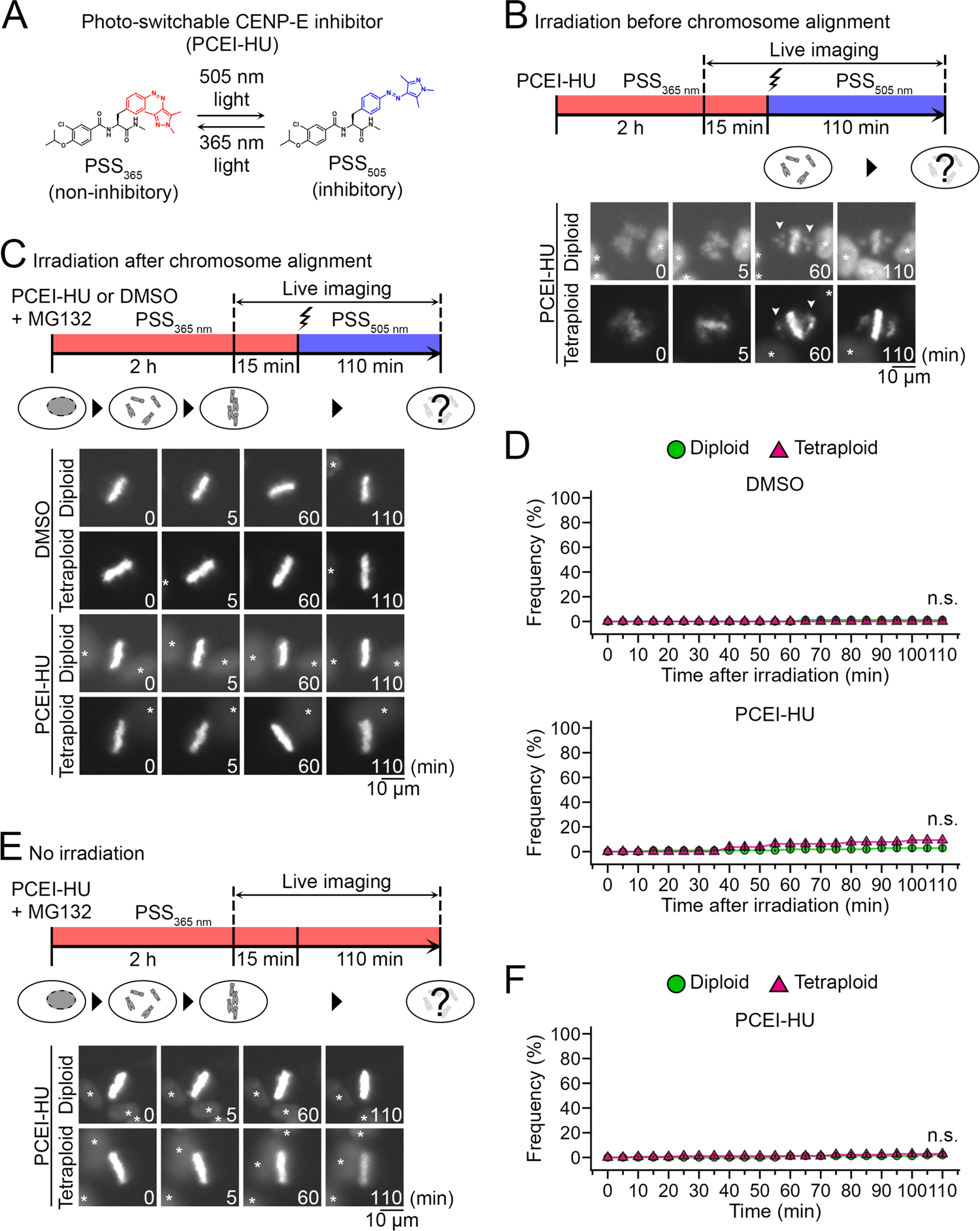
CENP-E inhibition does not impair the maintenance of the pre-aligned metaphase chromosomes. (**A**) Photoisomerization of the photo-switchable CENP-E inhibitor, PCEI-HU. (**B**, **C**, **E**) Schemes (top) and time-lapse images (bottom) of mitotic progression in HAP1 cells treated with DMSO or PCEI-HU. Cells were pre-treated with MG132 and SiR-DNA for blocking anaphase onset and staining chromosomes, respectively. Photo-switching of the inhibitor from the non-inhibitory PSS_365_to inhibitory PSS_505_ was induced before or after the completion of chromosome alignment in B or C, respectively. Note that the inhibitor blocked the equatorward movement of the misaligned polar chromosomes at PSS_505_ (B), whereas it did not affect the maintenance of the pre-aligned chromosomes (C). For comparison, we also tested chromosome movement in the cells treated with the inhibitor at PSS_365_ throughout the live imaging (E). Asterisks: Neighboring cells. (**D**, **F**) Cumulative frequency of de novo misalignment of the pre-aligned chromosomes in C or E (D or F, respectively). Mean ± SE of at least 44 cells from 3 independent experiments (n.s. between diploid and tetraploid cells at 110 min, the Brunner-Munzel test). Note that de novo misalignment was infrequent in diploids and tetraploids in all conditions.

**Fig. S7:**
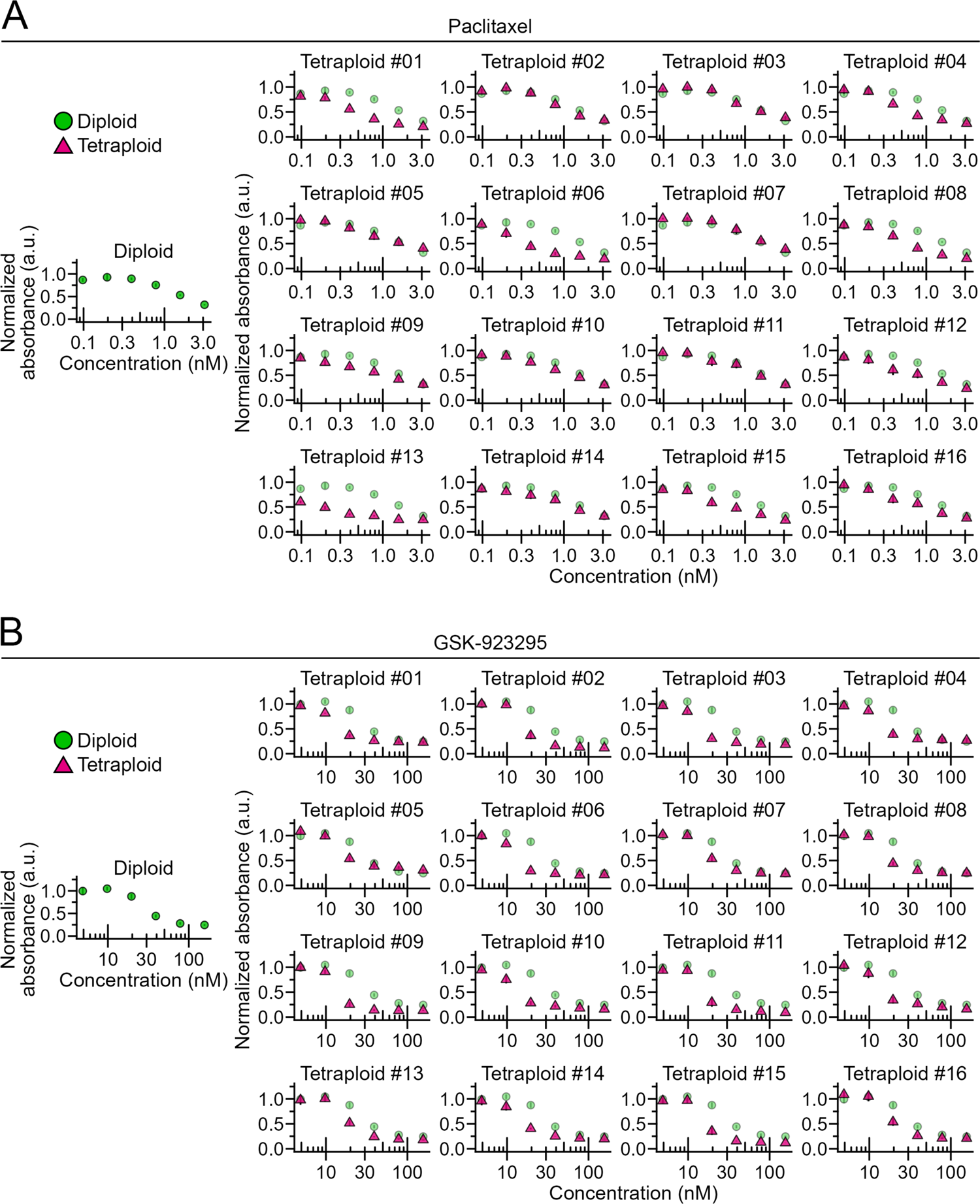
Proliferation of diploid or tetraploid HCT116 cells treated with different concentrations of paclitaxel or GSK-923295. (**A**, **B**) Dose-response curve of normalized absorbance in a comparative colorimetric cell proliferation assay using paclitaxel (A) or GSK-923295 (B) in diploid and tetraploid HCT116 cells. Mean ± SE of 4 samples from 2 independent experiments for each condition. For facilitating the comparison, identical dose-response plots of diploids were overlaid in all graphs of tetraploid plots.

